# Two *Stabiliser* loci in *Antirrhinum* suppress the transposition of Tam3 without compromising transposase production

**DOI:** 10.1101/2024.06.24.600476

**Authors:** Shasha Wang, Takako Uchiyama, Hiroyuki Kuwabara, Megumi Hirata, Ikumi Yuasa, Kenji Nakahara, Cathie Martin, Yuji Kishima

**Author notes:** Corresponding author: Yuji Kishima.

## Abstract

*Antirrhinum majus* has been known to exhibit occasional instabilities that are manifested as variegations and morphological chimeras. *Stabiliser* (*St*) is a historical locus that stabilizes phenotypically unstable or mutable traits in *Antirrhinum*. Here, we characterized two *St* loci, the previously described *Old Stabiliser* (*OSt*) and *New Stabiliser* (*NSt*), in *Antirrhinum* that specifically suppress the transposition of the Class II DNA transposable element, Tam3. Both *St* loci involve derivatives of Tam3 with unique structures: *OSt* has a pseudo-Tam3 copy whose 5’-terminal region has been rearranged compared to the cognate Tam3 element, and *NSt* consists of two intact copies of Tam3 in a head-to-head orientation. Neither locus interferes with the production of the intact Tam3 transposase (TPase) or the nuclear import of TPase.

Both *OSt* and *NSt* produce specific sRNAs from their 5’ terminal regions containing multiple TPase binding motifs. These specific sRNAs could repress Tam3 transposition by interacting with the TPase binding motifs within the Tam3 element or with the TPase itself.

## Introduction

The parasitism of transposable elements (TEs) as selfish genetic elements within their host organisms has been accompanied by evolution of a comprehensive set of master regulators (MRs) that control transposition of collective TEs in an epigenetic manner, inactivating the transcription of their genes and suppressing the production of the enzymes that catalyze transposition (Levin and Moran, 2011; Viviani et al., 2021; Liu et al., 2022). Consequently, few transcription or translation products are generally detected in hosts with inactivated TEs. However, some DNA transposons remain “surreptitiously” active even in repressive genomic environments. Examples of active plant DNA transposons are Ac/Ds, En/Spm, Mu in maize, Dart in rice, Tpn in morning glory, and Tam in *Antirrhinum* (Coen et al., 1989; Fedoroff et al., 1989; Inagaki et al., 1994; Gierl, 1996; Lisch, 2002; Fujino et al., 2005; Walbot and Rudenko, 2007; Lazarow et al., 2013). However, the frequency of excision of even active DNA TEs may be suppressed in response to specific host functions (Black et al., 1987; Lee et al., 1998; Slotkin et al., 2003, 2005).

Instability traits such as flower petal variegation often observed in *Antirrhinum majus* have been of interest since the 19th century. Petal variegation in *A. majus* occurring at the *Pallida* locus, which encodes dihydroflavonol 4-reductase (DFR) required for anthocyanin biosynthesis was observed by Darwin and De Vries and later referred to *palllida*^recurrens^ (*pal*^rec^) (Coen et al., 1989). Harrison and Fincham (1964) reported that an unlinked factor in *Antirrhinum* called *Stabiliser* (*St*) suppressed the petal variegation at *pal*^rec^. Petal variegation at the *Nivea* (*Niv*) locus encoding chalcone synthase responsible for the *niv*^recurrens^ (*niv*^rec^) allele and caused by Tam1 was not repressed by *St* (Harrison and Carpenter, 1973). However, a second unstable mutation in the *Nivea* locus caused by insertion of Tam3 (*niv*^rec^:98) was strongly repressed by *St* in a semi-dominant manner (Carpenter et al., 1987). Therefore, these studies led to suggestion that *St* is able to immobilize the transpositions of any copies of Tam3 *in trans*. An additional locus, similar phenotypically to *St* that can reduce the frequency of somatic excision of Tam3 copies was identified as having arisen spontaneously in an independent lineage of the *niv*^rec^:98 allele (Hashida et al., 2006). We renamed the historic *St* locus as *Old Stabiliser* (*OSt*) and the new locus *New Stabiliser* (*NSt*) with respect to their order of identification (Uchiyama et al., 2008).

Generally, transposition of Tam3 is not completely repressed in *A*. *majus*, but activity is regulated by temperature and is low temperature-dependent (LTDT) (Carpenter et al., 1987). The Tam3 transposase (Tam3 TPase) that catalyzes transposition is localized mainly at the plasma membrane at growth temperatures of 25°C (Fujino et al., 2011; Zhou et al., 2017). Therefore, Tam3 TPase is not present in the nucleus and cannot act on the Tam3 element in the nuclear DNA, resulting in the immobilization of Tam3 copies and the stable phenotypes associated with Tam3 unstable mutations at 25°C. Conversely at growing temperatures of 15°C, Tam3 TPase can enter to the nucleus and promote Tam3 transposition. The changes in subcellular localization of Tam3 TPase in response to temperature involved in LTDT are associated with the BED zinc finger motif (Znf-BED) of Tam3 TPase (Fujino et al., 2011; Zhou et al., 2017).

The *St* genes suppress Tam3 transposition at low temperature by mechanisms which seem to be unrelated to the processes of the TPase production (Uchiyama et al., 2008). This study addresses how the two *St* loci specifically suppress the Tam3 transposition without compromising normal Tam3 TPase function.

### Results Isolation of *OSt*

When *ost*/*niv*^rec^ (HAM2) was grown at 15 ℃, the petal lobes of its flowers showed strong variegation (> 1000 spots per flower) with a pale pink background due to Tam3 excision from the promoter in the *niv*^rec^ allele (Fig. 1A). PCR could detect somatic excision of Tam3 as a 2.0 kb band when *ost*/*niv*^rec^ was grown at 15 ℃ (Fig. 1A). Since Tam3 did not excise when grown at 25 ℃, no variegation of the *ost*/*niv*^rec^ petal appeared, and no excision band was detected (Fig. 1A). We confirmed that no PCR products reporting excisions of Tam3 from the wild type *Niv* allele were detectable (Supplementary Fig. 1). *OSt*/*niv*^rec^ (HAM21) shares the same *niv*^rec^ allele with *ost*/*niv*^rec^, while *OSt* is unique (Fig. 1A). Since Tam3 activity could be monitored by the *niv*^rec^ allele even at 15 °C, the weak variegation (< 20 spots per flower) observed in *OSt*/*niv*^rec^ suggested that *OSt* suppressed Tam3 transposition (Fig. 1A). PCR did not detect the 2.0 kb band for somatic excision of Tam3 from the promoter in the *niv*^rec^ allele of HAM21 when run for ≥ 30 cycles (Fig. 1A).

**Fig. 1.**
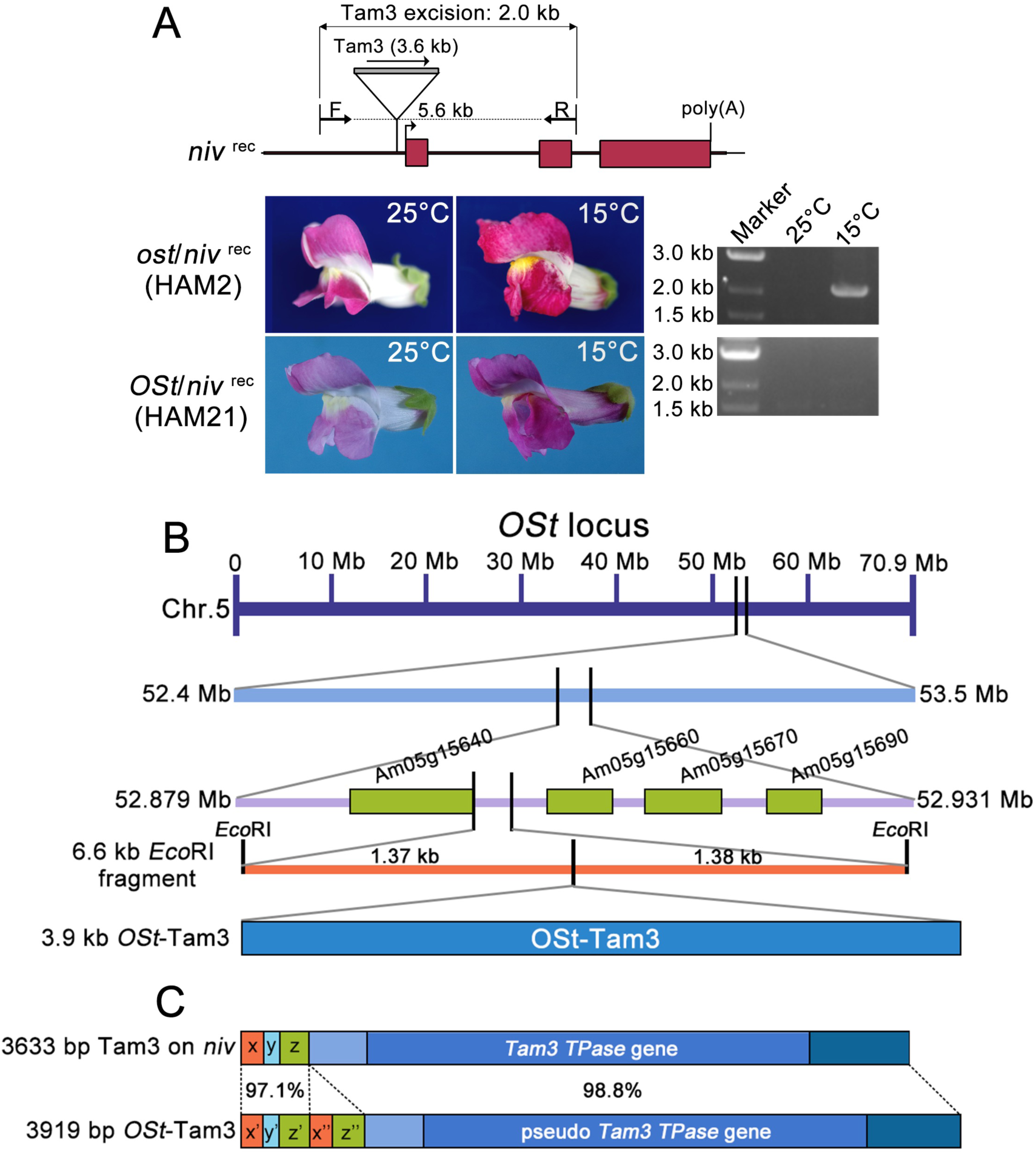
*OSt* suppresses Tam3 transposition. A. Profiles of *ost*/*niv*^rec^ (HAM2) and *OSt*/*niv*^rec^ (HAM21) *A. majus* lines. The two lines share the *niv*^rec^ allele at *Niv* locus encoding chalcone synthase (CHS), where a Tam3 element is inserted in the promoter of *CHS* gene. Tam3 insertion can restrict the expression of *CHS* gene in *niv*^rec^, which thus gives rise to the flower petals with pale red background at 25℃. *ost*/*niv*^rec^ (HAM2) does not contain suppressors for Tam3 transposition and thus exhibits red sites and sectors on flower petals at 15℃ due to Tam3 transposition. *OSt* suppressor is carried in the *OSt*/*niv*^rec^ (HAM21) genome and strongly suppresses Tam3 transposition, which eventually causes a few red sectors on flower petals at 15℃. Additionally, PCR analysis using F and R primers to examine Tam3 transposition showed that Tam3 is inactive at 25℃, and 2.0 kb Tam3-excision bands were missing in *ost*/*niv*^rec^ (HAM2) and *OSt*/*niv*^rec^ (HAM21). At 15℃, Tam3 is active in *ost*/*niv*^rec^ (HAM2), and the excision bands can be observed, but not in *OSt*/*niv*^rec^ (HAM21). B. The chromosomal position of *OSt*. C. Structural comparison between Tam3 on *niv*^rec^ and *OSt*-Tam3.

To confirm that *OSt* is single semi-dominant locus as described by Harrison & Fincham (Harrison and Fincham, 1968), we crossed *ost*/*niv*^rec^ (HAM2) ×*OSt*/*niv*^rec^ (HAM21). The F_2_ population segregated into 118 high- and middle-spotted plants (> 100 spots per flower: we did not discriminate between high and middle spots in this population) and 35 low-spotted plants (< 20 spots per flower) (Supplementary Table 1). An independent cross was performed between *OSt*/*niv*^rec^/*Pal* (HAM21) and *ost*/*Niv*/*pal*^rec^ (HAM22), and we selected the 51 plants possessing either homozygous *niv*^rec^ and/or homozygous *pal*^rec^ among the F_2_ plants. These 51 plants segregated into 9 high-(> 500 spots per flower), 31 middle-(> 50 spots per flower) and 11 low- (< 20 spots per flower) spotted plants in the F_2_ population (Supplementary Table 1). These segregations in the F_2_ populations of the crosses of *OSt*/*niv*^rec^ × *ost*/*niv*^rec^ and *OSt*/*niv*^rec^/*Pal* × *ost*/*Niv*/*pal*^rec^ statistically fitted the ratios of 3:1 and 1:2:1, respectively (Supplementary Table 1 and Supplementary Fig. 2A). These segregation results supported the hypothesis that the low-spotted *niv*^rec^ phenotype is regulated by a single semi-dominant locus, *OSt*.

Several reports have suggested that active DNA transposons can be silenced by their own, transposon-derived suppressors, and that such suppressors are produced by transposon recombination, such as *KP* of the P element in *Drosophila* (Black et al., 1987) and *Mu killer* (*Muk*) of the MuDR partial element in maize (Slotkin et al., 2003). Thus, *OSt* might have been derived from alterations to a Tam3 element arising from recombination. Supporting such an idea, Harrison & Fincham (1967) reported that they observed a loss of *OSt* function somatically, in the form of higher frequency of sectors on low spotted *pal^rec^* petals. To explore this possibility, transposon tagging was performed to investigate whether a specific Tam3-associated fragment exists in the genome of *OSt* plants using the ORF region of Tam3 as a probe.

We obtained a 6663 bp Tam3-associated DNA restriction enzyme fragment as a candidate tag for the *OSt* locus following a transposon tagging procedure (Fig. 1B and Supplementary Description 1). Within this, a 3919 bp candidate *OSt-*Tam3 element was located on chromosome 5 (Supplementary Fig. 3). *OSt*-Tam3 encoded a pseudo-*Tam3 TPase* gene with 98.8% homology to the intact element cloned from the *Niv* locus from line *niv*^rec^:98 (Hehl et al., 1991). Notably, the 5’ region of the 3919 bp *OSt*-Tam3 element contained an extra 286 bp sequence caused by rearrangements of the 5’ end of Tam3 (Fig. 1C and Supplementary Fig. 4A). In the 5’ region of the reference Tam3 element, the region 1-376 bp can be divided into three parts, x-y-z.

The corresponding region consisting of x’-y’-z’ in *OSt*-Tam3 showed 97.1% to the reference element (Fig. 1C). Dot plot analysis indicated that regions x’ and z’ had been duplicated in *OSt*-Tam3, resulting in new sequence consisting of five parts, x’-y’-z’-x’’-z’’ (Fig. 1C and Supplementary Fig. 5A). There are 11 Tam3 TPase binding GCHCG motifs in the x region of Tam3, and they are also conserved in x’ and x’’ regions of *OSt*-Tam3 (Supplementary Fig. 5B). Compared with the 5’ region in intact Tam3, the corresponding parts in *OSt*-Tam3 also showed several single nucleotide polymorphisms (SNPs) (Supplementary Fig. 5B). Either this sequence rearrangement and/or SNPs found in *OSt*-Tam3 might be the cause of the suppression of Tam3 transposition associated with the *OSt-*Tam3 locus.

### Isolation of *NSt*

*NSt*/*niv*^rec: beni^ (HAM3) was derived from *nst*/*niv*^rec: beni^ (HAM5) and was identified as a spontaneous mutant of *NSt* (Hashida et al., 2006; Uchiyama et al., 2008). *nst*/*niv*^rec: beni^ shows red spots and sectors on the white background petal (> 1000 spots per flower) at 15℃ due to Tam3 transposition from the *niv*^rec: beni^ allele (Uchiyama et al., 2013) (Fig. 2A). Both the lines share the *niv*^rec: beni^ allele, whereas *NSt*/*niv*^rec: beni^ shows no variegation (<10 spots per flower) at 15℃, indicating immobilization of Tam3 at the *niv*^rec: beni^ locus (Fig. 2A). PCR detected the 2.0 kb Tam3 excision band from *niv*^rec: beni^ in *nst*/*niv*^rec: beni^ grown at 15℃, but not in *NSt/niv*^rec: beni^ or the plants grown at 25℃, confirming that *NSt/niv*^rec: beni^ carries the *NSt* locus that suppresses Tam3 transposition (Fig. 2A).

**Fig. 2.**
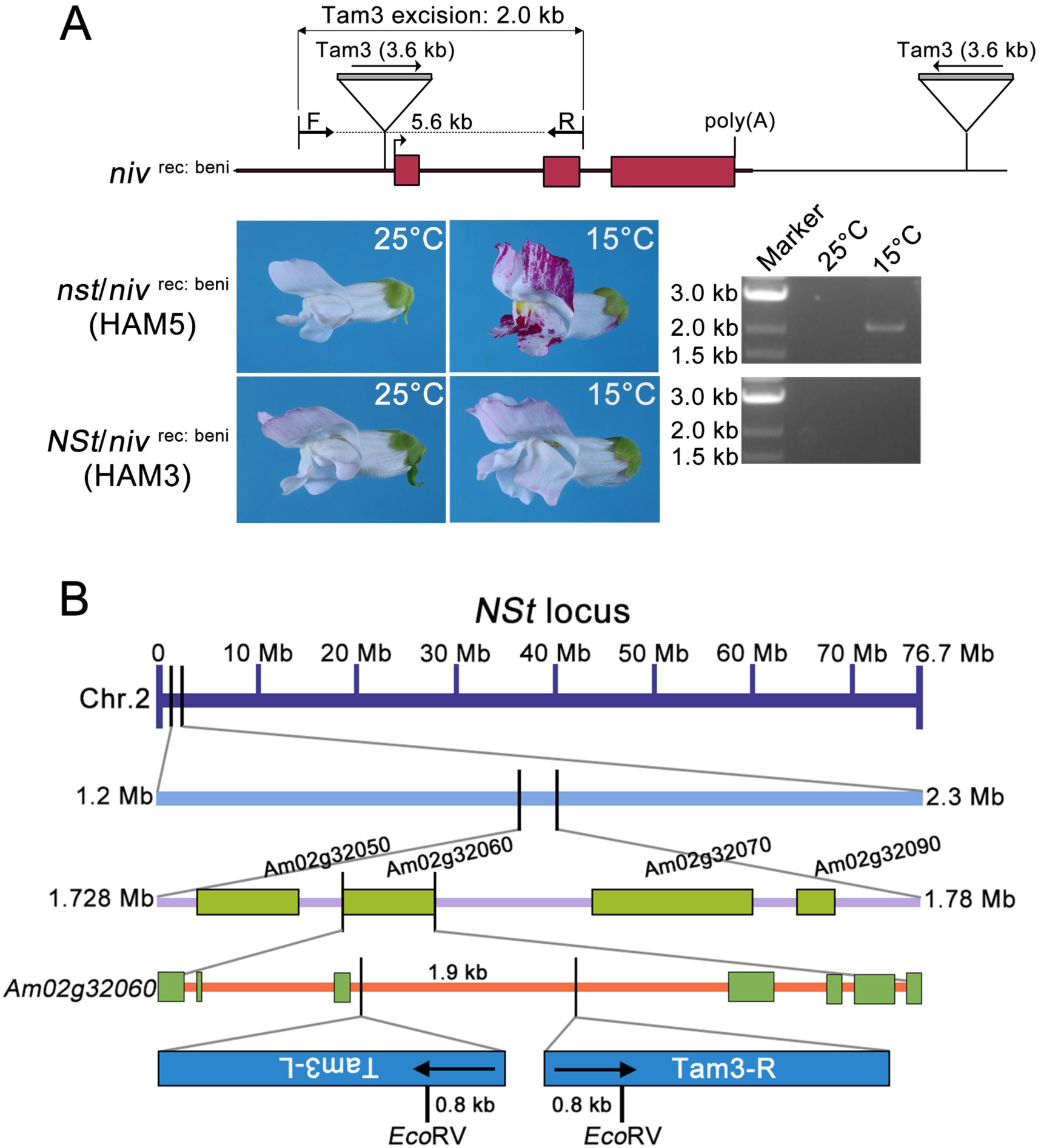
*NSt* suppresses Tam3 transposition. A. Profiles of *nst*/*niv*^rec: beni^ (HAM5) and *NSt*/*niv*^rec: beni^ (HAM3) *A. majus* lines. The two lines share the *niv^rec:^* ^beni^ allele at *Niv* locus, where two Tam3 elements are inserted into the promoter and downstream of *CHS* gene, respectively. Tam3 strongly restricts the expression of *CHS* gene at *niv*^rec: beni^ allele, which causes light-colored flower petals at 25℃. *nst*/*niv*^rec: beni^ (HAM5) does not contain suppressors for Tam3 transposition and thus exhibits red sites and sectors on flower petals at 15℃ due to Tam3 transposition. *NSt* suppressor resides in *NSt*/*niv*^rec: beni^ (HAM3) genome and strongly suppress Tam3 transposition, which results in no variegations on flower petals at 15℃. The 2.0 kb Tam3-excision band was not observed in the two lines at 25℃ and *NSt*/*niv*^rec: beni^ (HAM3) at 15℃, but was observed in *nst*/*niv*^rec: beni^ (HAM5) at 15℃. B. The chromosomal position and structure of *NSt*.

We crossed *NSt/niv*^rec: beni^ (HAM3) × *nst/niv*^rec: beni^ (HAM5) twice, independently (Supplementary Fig. 2B). A total of the 214 F_2_ progenies of *NSt*/*niv*^rec: beni^ × *nst*/*niv*^rec: beni^ segregated into two different classes: high-spots (> 1000 spots per flower), and low-spots (<10 spot per flower) with a 1: 3 ratio of the number of plants (Supplementary Table 1 and Supplementary Fig. 2B). These results confirmed that excision of Tam3 from the *niv*^rec: beni^ locus was controlled by a single dominant locus (*NSt*) as reported by Hashida *et al*. (2006).

By analogy to the method used for identification of *OSt*, we undertook transposon tagging using Tam3 as a probe to identify the Tam3-associated fragment specifically linked to the *NSt* phenotype. Detailed procedures for transposon tagging of *NSt* are described in Supplementary Description 2. We identified an *Eco*RV restriction enzyme DNA fragment linked to *NSt*, which was 3.5 kb long and contained 0.8 kb of Tam3 sequences at the two ends of the *Eco*RV fragment with a 1.9 kb internal sequence (Supplementary Fig. 6). Further structural analysis of the sequence surrounding the 3.5 kb fragment in the *NSt*/*niv*^rec: beni^ (HAM3) genome revealed two intact Tam3 sequences in head-to-head orientation inserted into the third intron of the *Am02g32060* gene encoding transcription factor BREVIS RADIX protein (Fig. 2B). Dot plot analysis of the sequence in this region showed that the two Tam3 sequences have the same structure as the reference element in the *niv*^rec^ allele (Hehl et al., 1991) (Supplementary Fig. 7A). In the *nst*/*niv*^rec: beni^ (HAM5) genome, the right-hand copy of the two Tam3 sequences is absent (Supplementary Fig. 7B). Hence, the *NSt* locus of *NSt*/*niv*^rec: beni^ (HAM3) may have arisen by *de novo* insertion of Tam3 on the right side of the Tam3 element in the *nst* allele of *nst*/*niv*^rec: beni^ (HAM5) (Supplementary Fig. 7B). The *Am02g32060* gene residing *NSt* was found in the Snapdragon genome database to be located at the end of chromosome 2 (Fig. 2B and Supplementary Fig. 3).

### Transcription, translation, nuclear import, and DNA-binding of Tam3 TPase

We then examined how the two *Stabiliser* loci inhibit Tam3 transposition. Tam3 transposition requires four steps involving the *Tam3 TPase* gene: transcription, translation, nuclear import, and binding to Tam3 elements. Regarding the transcription and translation of the *Tam3 TPase* gene, we performed qRT-PCR to detect differences in *Tam3 TPase* gene expression and immunoblot analysis to examine the translated product in the *OSt* (HAM21), *NSt* (HAM3), and *ost*/*nst* (HAM5) (Supplementary Description 3). The transcription and translation for the *Tam3 TPase* gene could not be quantitatively distinguished between *OSt*, *NSt*, and *ost*/*nst* plants (Supplementary Fig. 8; Supplementary Description 3). These results confirmed previous reports that *OSt* and *NSt* do not suppress the *Tam3 TPase* gene (Uchiyama et al., 2008).

We then examined the nuclear import of Tam3 TPase in the presence of *OSt* and *NSt*. When plants are grown at 15℃, Tam3 TPase is synthesized in the cytoplasm and enters the nucleus. We tested the nuclear import of Tam3 TPase by transient assay using *Antirrhinum* protoplasts transformed with a construct for GFP-fused Tam3 TPase in the presence of *OSt* and *NSt.* We found that nuclear-localized GFP signals were observed more frequently in all protoplasts from *OSt*, *NSt*, or *ost*/*nst* plants at 15℃ compared to 25℃, indicating that nuclear import of Tam3 TPase at the low temperatures occurs in *OSt* and *NSt* genetic backgrounds, and confirming early reports that temperature and *OSt* influence Tam3 transposition independently (Fig. 3A and B; Supplementary Table 2) (Fincham and Harrison, 1967).

**Fig. 3.**
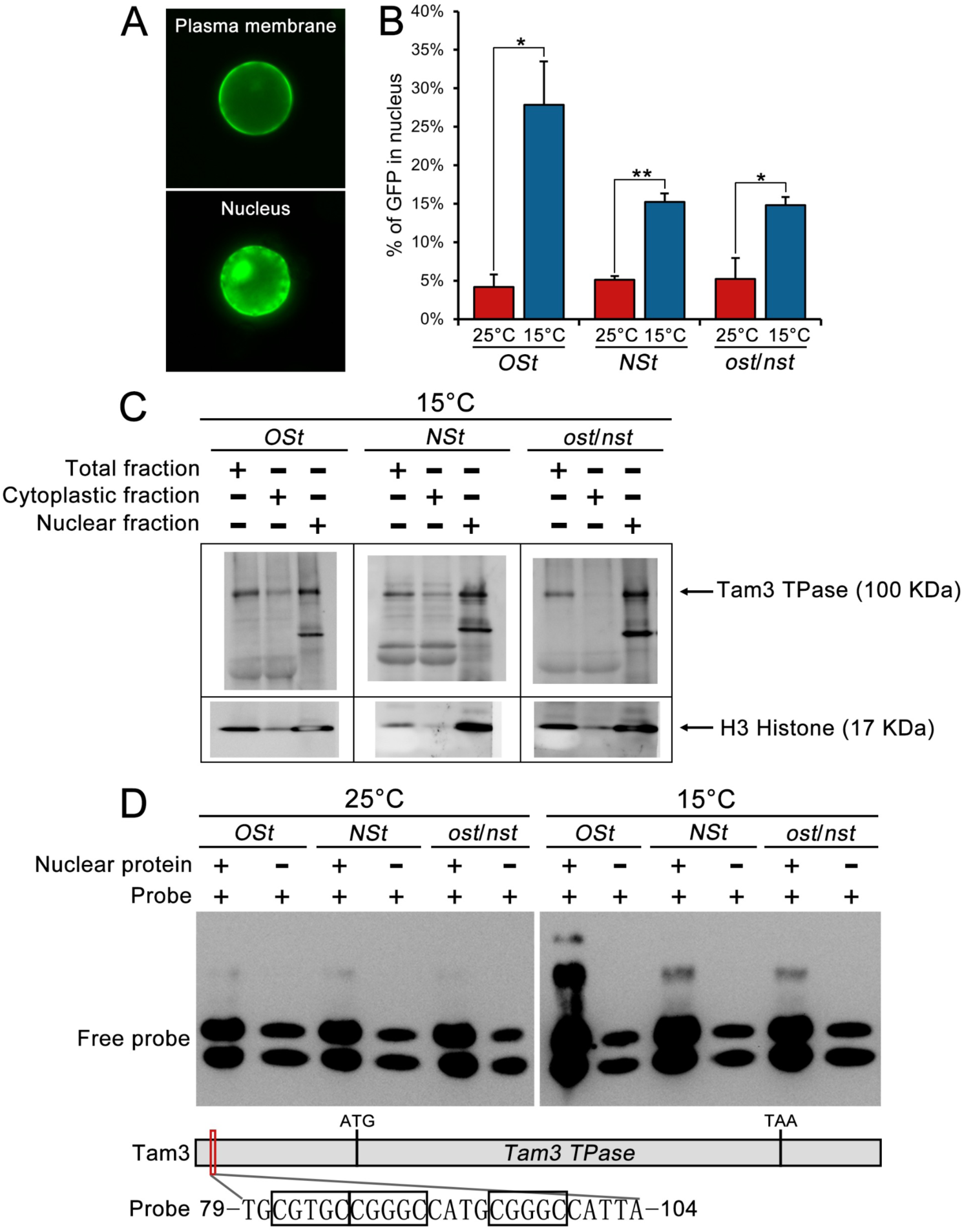
Nuclear import and Tam3 DNA binding of Tam3 TPase. A. Plasma membrane and nuclear localization of Tam3 TPase in *Antirrhinum* protoplasts. B. The proportions of cells with Tam3 TPase-GFP signals in nuclei out of total number of cells with fluorescence (Supplementary Table 2). Data represent means ± SD. Significant differences were determined by Student’s t-test. C. Immunoblot analysis of total, cytoplastic and nuclear protein. The fragments indicated by upper and lower arrows correspond to 100 kDa of Tam3 TPase and 17 kDa of H3 Histone, respectively. The plus (+) and minus (-) marks represent the presence and absence of each protein in each lane, respectively. D. EMSA analysis of DNA binding ability of nuclear Tam3 TPase. Nucleotide sequence of Tam3: 79-104 in Tam3 containing three GCHCG motifs, was used as a probe. The probe and nuclear protein are mixed with 50 fM and 2 ug, respectively. Shifted bands from free probes (bottom two bands) were detected in *OSt*, *NSt* and *ost*/*nst* plants grown at 15℃ when nuclear protein was added (+). While absence of nuclear protein (−), no shifted bands appeared.

Following these experiments, we concluded that the two *Stabilisers* are probably involved in the final step of Tam3 transposition, the binding of Tam3 TPase to Tam3 DNA sequences, to limit Tam3 transposition. We therefore investigated the effect of *OSt* and *NSt* on the binding affinity of Tam3 TPase to its target sites. Electrophoretic mobility shift assay (EMSA) was performed using a 26 bp fragment containing three Tam3 TPase target motifs (GCHCG) as a probe (Hashida et al., 2006). Nuclear protein extracted from young leaves was employed to test the interaction between Tam3 TPase and the 26 bp target fragment. We confirmed that TPase protein extracted from all the plants grown at 15 ℃ was detected as probe-protein shift bands in nuclear fractions (Fig. 3C and D). Interestingly, strongly shifted bands were observed in the extracts from *OSt* plants grown at 15 ℃, which was consistent with the higher frequency of nuclear localized GFP signals in the protoplasts (Fig. 3B). However, neither *OSt* and *NSt* altered the binding affinity of Tam3 TPase to its target motifs.

### mRNA profiles in *OSt* and *NSt*

According our results and those of others (Chatterjee and Martin, 1997), it was assumed that *OSt* and *NSt* interfere with the ability of the Tam3 TPase to bind to its target sequences on the chromosome, without compromising Tam3 TPase production. Firstly, we investigated whether the *OSt*- and *NSt-* factors might be RNA products, so we carried out poly(A)-enriched RNA-seq analysis to explore the transcription of the *Tam3 TPase* gene in mRNA from the *OSt*, *NSt*, and *ost*/*nst* plants. Similar transcriptional patterns of the sense strand of *Tam3 TPase* open reading frame (ORF) were observed in these three genotypes, confirming the results of qRT-PCR and immunoblotting analyses (Supplementary Figs. 8-9). In *OSt* plants we found mRNA transcripts in the region of the pseudo-*Tam3 TPase* gene, although no transcripts were detected covering the 5’ rearranged region of *OSt*-Tam3 (Supplementary Fig. 10). In *NSt* plants mRNA transcripts were observed for the internal region between two Tam3 elements in *NSt* locus, corresponding to transcripts from the third intron of the *Am02g32060* gene, but these aberrant transcripts did not align with the TPase transcript and therefore were unlikely to interact directly with the TPase (Supplementary Fig. 11). Although the two *Stabiliser* genes, *OSt* and *NSt*, gave rise to different mRNA expression patterns between the three genotypes, these different patterns of the RNA-seq data could not be related directly to the suppression of Tam3 transposition of *OSt* and *NSt*.

### lncRNA profiles in *OSt* and *NSt*

The mRNA patterns in *OSt* and *NSt* did not show that high-expression poly(A)-tailed mRNAs were responsible for the suppression of Tam3 transposition. Thus, we investigated whether other RNAs, non-coding RNAs, might be derived from *OSt* and *NSt* loci and inhibit the binding of Tam3 TPase to Tam3 DNA sequences. One type of non-coding RNA is long non-coding RNA (lncRNA) the length of which is longer than 200 nucleotides and has important biological functions, including interacting with proteins to inhibit their binding to specific regions of DNA and interacting with DNA to decrease chromatin accessibility (Statello et al., 2021). Therefore, we performed rRNA-depleted RNA-seq analysis to investigate whether lncRNAs were generated from *OSt* and *NSt* loci. Within the Tam3 element, lncRNA transcripts were found in the *Tam3 TPase* gene in *OSt*, *NSt* and *ost*/*nst* plants (expression level: *OSt* > *NSt* = *ost*/*nst*), but no lncRNA transcripts were detected in both the terminal GCHCG-enriched regions of Tam3 (Supplementary Fig. 12). In *OSt*, we found lncRNA transcripts located in the pseudo-*Tam3 TPase* gene, although transcripts were missing for the 5’ rearranged region of *OSt*-Tam3 (Supplementary Fig. 13). In the *NSt* locus, abundant lncRNA transcripts were detected in the inter-element region between Tam3-L and -R in *NSt* plant, and a few transcripts existed in the same region in *ost*/*nst* plant (Supplementary Fig. 14). However, we were unable to infer any direct relationship between *NSt*-derived lncRNAs and Tam3 inactivation. Therefore, the results of different RNA expression patterns in the three plant lines could not be directly associated with the Tam3 silencing caused by either of the two *Stabiliser* loci.

### sRNA profiles of Tam3 in *OSt* and *NSt*

Based on knowledge of sRNAs that play a role in transposon inactivation (Castel and Martienssen, 2013; Ito, 2013), we performed sRNA-seq analyses to investigate whether sRNAs are associated with Tam3 inactivation in either *OSt* or *NSt* plants. The functions of sRNAs are generally determined by their size: 21 nt and 22 nt for mRNA cleavage and 24 nt for DNA methylation (Kryovrysanaki et al., 2022). sRNAs from *OSt*, *NSt* and *ost/nst* plants were classified by function and compared between the three genotypes in terms of their distribution and abundance (Supplementary Figs. 15-16). 24 nt sRNAs on both sense and antisense strands of the Tam3 sequence accumulated more than the other sRNAs from the three genotypes. sRNAs were detected mainly in the coding region of the TPase (690 - 3101 bp of the Tam3 element), the distribution patterns of which were more or less similar between the three genotypes (Supplementary Figs. 15-16). We found no clear difference in either the accumulation or distribution patterns of sRNAs between the three genotypes. However, despite the accumulation of sRNAs in the 5’ terminal region of the Tam3 element being low, the distribution patterns were different between the three genotypes (Supplementary Figs. 15-16). We focused our investigations on the sRNAs from the 5’ terminal regions where Tam3 TPase binding motifs are present.

sRNAs from the x and y regions of the 5’ terminus of Tam3 could be associated with the *Stabiliser* properties (Fig. 4), while, in contrast, the accumulations of sRNAs were detected only weakly in the 3’ terminus of the three genotypes (Supplementary Fig. 17). The three genotypes differentially accumulated sRNAs from the sense strand of the x and y regions of Tam3 (Fig. 4), suggesting that these sRNAs may bind to the antisense strand. In *OSt* plants, sRNAs from the first 30 bp region of the 5’ terminus of Tam3, which contains the terminal inverted repeat (TIR) sequence and a GCHCG motif on the antisense strand, selectively accumulated (Fig. 4). These sRNAs included those with a single SNP on the 13th nucleotide position from the 5’ end of the x’ and x’’ segments in *OSt*-Tam3 (Supplementary Fig. 5B). In *NSt* plants, sRNAs from the sense strand of the 70-150 bp region accumulated strongly and might protect binding to the x segment containing four TPase GCHCG binding motifs corresponding to the antisense sequence (Fig. 4). Compared with *OSt* and *NSt*, sRNAs from *ost*/*nst* plants were not observed at the detectable levels from the sense strand sequence (Fig. 4). For the antisense strands, a relatively low accumulation of sRNA from *OSt* and *NSt* was observed for the 30-70 bp region of the x segment which includes six GCHCG motifs corresponding to the sense strand sequence (Fig. 4).

**Fig. 4.**
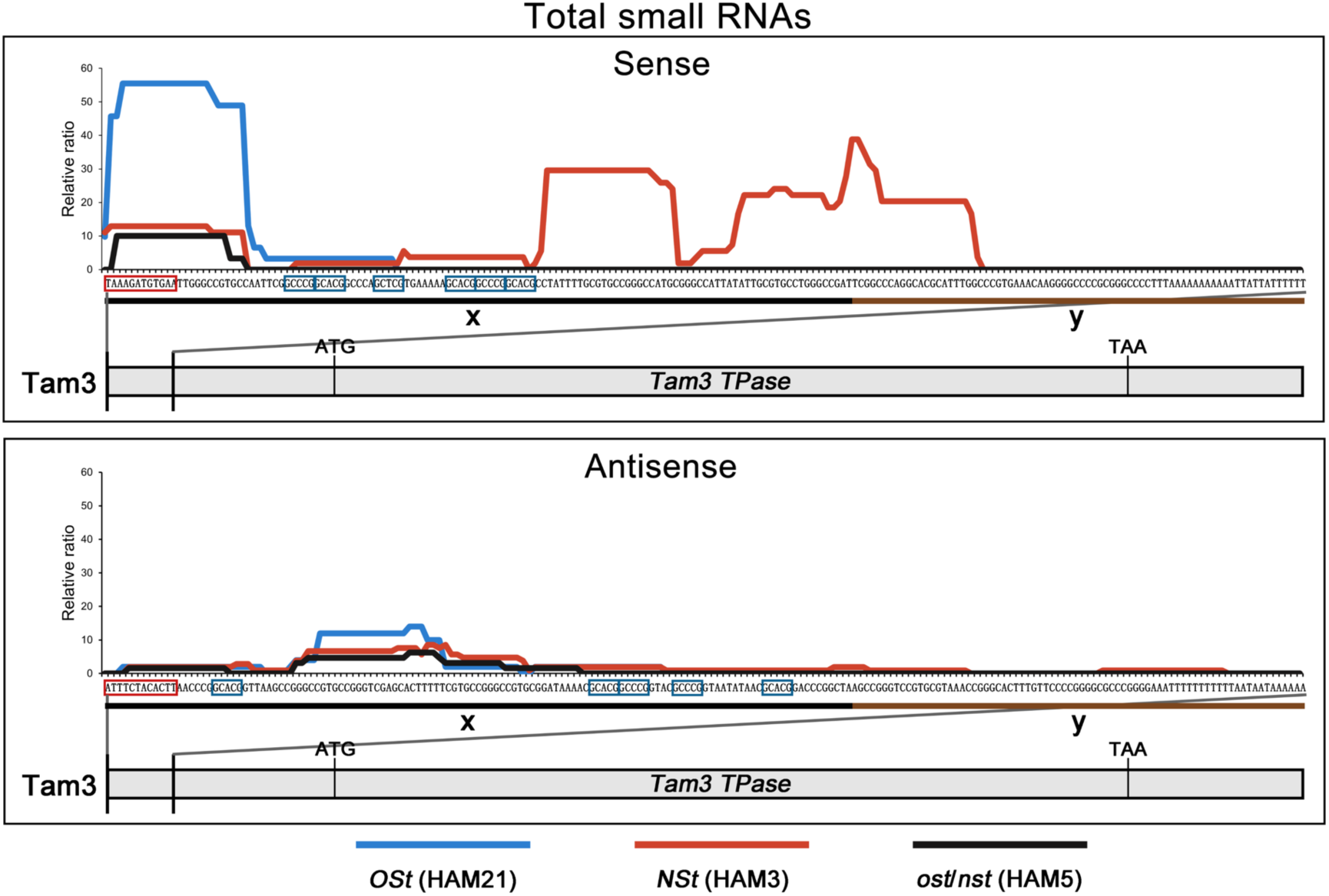
Distributions and relative amounts of total small RNAs in 5’ region of intact Tam3. Except for the *OSt* line, the sRNAs that were perfectly matched to the 5’ region of Tam3 (1-201 bp) were analyzed in *NSt* and *ost*/*nst* lines. In the *OSt* line sRNAs that contain zero and one SNP were aligned to 5’ 1-201 bp region of Tam3. The red and blue squares on nucleotide sequences indicate Terminal Inverted Repeat (TIR) and GCHCG motifs, respectively. The lines above the DNA sequence indicate as follows; blue: *OSt* (HAM21), red: *NSt* (HAM3), black: *ost*/*nst* (HAM5). Black and brown lines below the DNA sequence represent “x” and “y” regions of intact Tam3, respectively. Relative ratios were calculated as follows; the RPH (reads per hundred mapped reads) of each line divided by the average RPH of *ost*/*nst* line within 5’ 1-201 bp region of Tam3.

### sRNA profiles in *OSt* and *NSt* sequences

Based on the previous sRNA-seq results, we observed that specific sRNAs accumulated in the 5’ region of Tam3 sequence in *OSt* and *NSt* lines, compared to the *ost*/*nst* line (Fig. 4; Supplementary Figs. 15-16). In both the sense and antisense strands of the *OSt* sequence, most of the sRNAs were distributed in the region from the 5’ end of *OSt*-Tam3 to the end of the pseudo-*Tam3 TPase* gene (1-3385 bp of the *OSt*-Tam3 sequence) (Supplementary Fig. 18). In the 5’ end of the *OSt*-Tam3 sequence, 24 nt sRNAs came predominantly from the x’, x’’ and z’’ regions on the sense strand and from the z’’ region on the antisense strand (Supplementary Figs. 18 and 19A-B). We could detect sRNAs specifically from the antisense strand of the 5’ end of Tam3-L in the *NSt* allele, which was not the case for the *nst* allele (Supplementary Fig. 19C-D).

In general, double-stranded RNAs produced by pairing sense and antisense transcripts of DNA fragments and transcribing inverted repeats can be processed into sRNAs (Mathieu and Bender, 2004). The production of sRNAs on *OSt* and *NSt* sequences is most likely due to the bidirectional transcription of *OSt*-Tam3 and the transcription of two inverted Tam3 elements for *NSt*, respectively. We performed a promoter prediction on the 6675 bp surrounding the *OSt*-Tam3 sequence and the 9133 bp surrounding the *NSt* elements and mined for possible transcription start sites (TSSs) on the sense and antisense strands of the two sequences (Supplementary Table 3).

According to the TSS score, 32 sites on the sense and antisense strands of *OSt* and 46 sites on the sense and antisense strands of *NSt* were identified (Supplementary Table 3). Based on the distribution of the sRNAs and TSS scores in the *OSt*-Tam3, two TSS combinations were identified in the flanking and inside sequences (Fig. 5A). In *NSt*, since no antisense RNA was detected from the internal sequence between the two Tam3 elements (Supplementary Fig. 11 and 14), it is likely that only sense strand transcription occurred from the Tam3 on the left to the Tam3 on the right-hand side. This transcript could result in a hairpin structure with both Tam3 5’ sequences, giving rise to the specific sRNAs observed (Fig. 5B). Transcription from *OSt* and *NSt* could generate sense-antisense paired and hairpin (stem-loop) dsRNAs, respectively (Fig. 5). The predicted promoter activities on *OSt* and *NSt* alleles suggested that their respective transcription might generate sRNAs as Tam3-specific repressors.

**Fig. 5.**
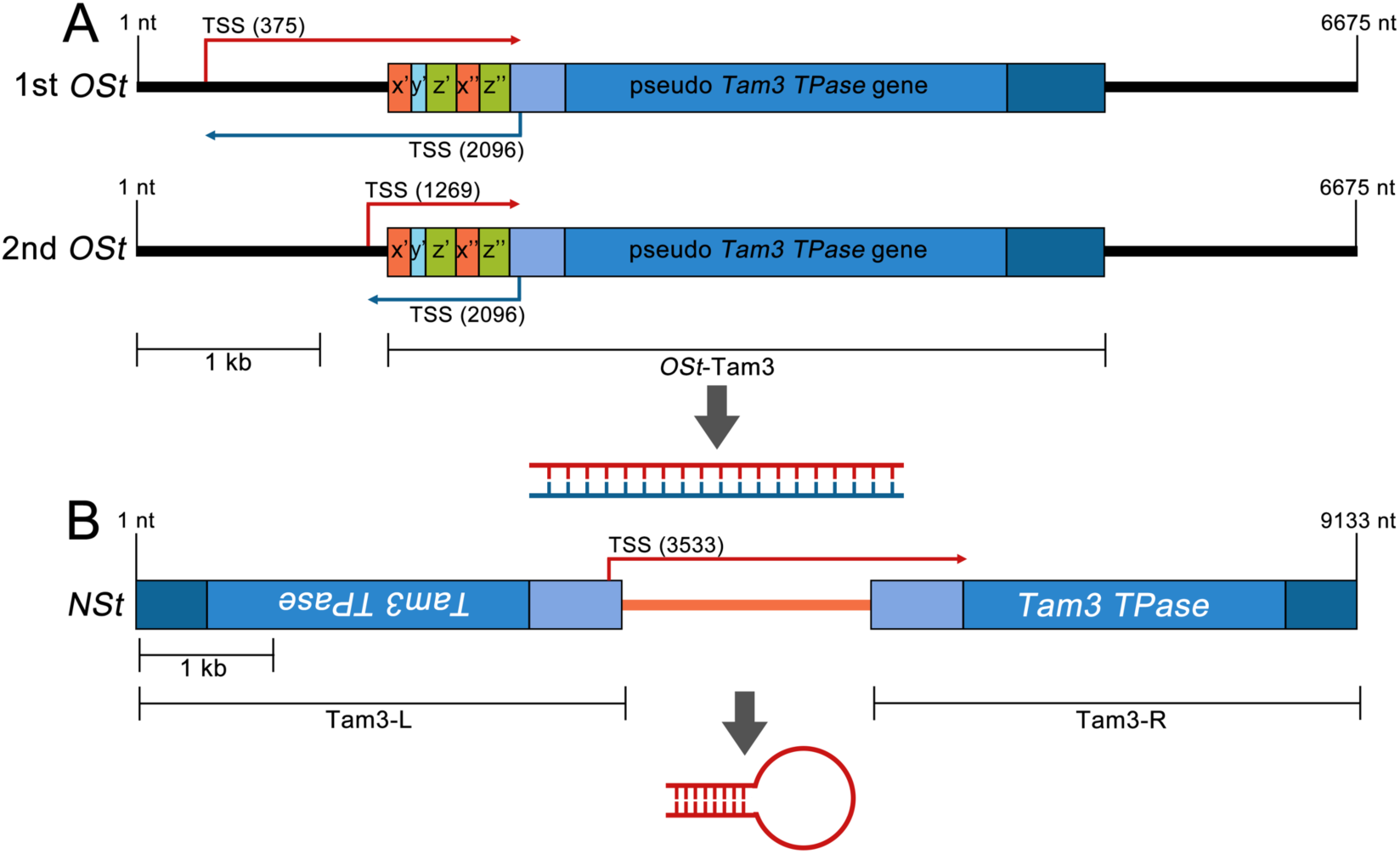
Possible double-stranded RNAs derived from *OSt* and *NSt*. A. Possible double-stranded RNAs derived from *OSt*. The two highest combinations of sense and antisense transcriptional start sites were represented in *OSt* (Supplementary Table 3). The transcription on the sense and antisense strands of *OSt* DNA fragment causes the production of sense and antisense transcripts which are paired to form double-stranded RNA. B. Possible double-stranded RNAs derived from *NSt*. One sense transcriptional start site was represented in *NSt*. The transcription starts from the 5’ end of Tam3-L and goes through to the 5’ end of Tam3-R. The transcript contains two paired parts at the two ends and thus can be folded into stem-loop structure where dsRNA exists in stem part. Red and blue vertical fold arrows represent predicted transcriptional start sites on sense and antisense strands of *OSt* and *NSt*, respectively. The number in parenthesis represents the nucleotide position of TSS from the left side of *OSt* or *NSt*. TSS, transcriptional start site.

### Detection of hairpin transcript in *NSt*

According to our finding of sense transcripts in the internal region between two Tam3 of *NSt*, it is most likely that the transcription of *NSt* from Tam3-L to Tam3-R could result in a hairpin structure which lie in head-to-head orientation. This hairpin transcript could be processed into the specific sRNAs. To examine for presence of the hairpin transcript, PCR analyses were performed for the stem and loop parts of the hairpin product using two pairs of primers, respectively (Supplementary Fig. 20). The two parts only can be detected in *NSt* hairpin transcript together (Supplementary Fig. 20). We were unable to amplify cDNAs beyond both Tam3 sequences, which may be because DNA polymerase was unable to extend to the double-stranded stem region of a stem-loop structure from the single-stranded loop (Zhong et al., 2019). The results suggested that transcription of *NSt* can result in a hairpin transcript with the 5’ end regions of Tam3 then responsible for the generation of the specific sRNAs.

## Discussion

Although the two *Stabiliser* loci have distinctive structures and chromosomal positions, both play suppressive roles in Tam3 transposition. The *OSt* locus, carrying the pseudo-Tam3 lies in an intergenic region on chromosome 5 of the *OSt*/*niv*^rec^ (HAM21) line. The unique feature of the *OSt* locus is that its causative rearrangement occurred in the terminal region of the element, outside of the *TPase* coding sequence. The *NSt* locus on chromosome 2 involves two Tam3 fragments inserted in head-to-head orientation in the third intron of the *Am02g32060* gene in the *NSt*/*niv*^rec: beni^ (HAM3) line.

Currently, four TE-specific repressor genes have been identified such as *En-I102* for En/Spm in maize, *Muk* for Mu element in maize, and *KP* and *Lk-P(1A)* for P element in *Drosophila* (Black et al., 1987; Cuypers et al., 1988; Ronsseray et al., 1991; Slotkin et al., 2003). *En-I102* and *KP* are the deletion derivatives of En and P elements, respectively (Black et al., 1987; Cuypers et al., 1988). *Lk-P(1A)* and *Muk* share the head-to-head structure of their respective TEs (Roche and Rio, 1998; Slotkin et al., 2005). *Lk-P(1A)* has two intact P elements, whereas *Muk* was generated by duplication of partially deleted *MuDR* element (Ronsseray et al., 1996; Roche and Rio, 1998; Slotkin et al., 2005).

Based on the mechanisms of suppression, the four examples of DNA transposable element suppression can be divided into two groups: pre-translational suppression and post-translational suppression. The pre-translational repression group includes *Muk* and *Lk-P(1A)*, which generate small interference RNAs (siRNAs) for the degradation of *TPase* transcripts (Slotkin et al., 2005; Brennecke et al., 2008; Majumdar and Rio, 2015). These TE-specific repressor genes have a rearranged segment of the authentic TE structure, which could lead to siRNAs against the *TPase* gene transcript (Slotkin et al., 2005; Brennecke et al., 2008; Majumdar and Rio, 2015). The other two examples, *En-I102* and *KP*, involve post-translational suppression (Cuypers et al., 1988; Lee et al., 1998). Both *En-I102* and *KP* can produce aberrant TPase to compete with normal TPase for target sites on TEs (Cuypers et al., 1988; Lee et al., 1998).

*OSt* and *NSt* belong to a third group, with mechanisms distinct from pre-translational suppression or post-translational repression. These *Stabiliser* genes do not restrict transcription and translation of the *TPase* gene like *Muk* and *Lk-P*(*1A*), nor do they generate aberrant truncated TPase to compete for target sites at the termini of the element, like *En-I102* and *KP*. All these results suggest that the mechanism(s) of repression by *OSt* and *NSt* on Tam3 transposition are distinct from the general epigenetic regulation of TEs (Wang et al., 2022).

The *OSt* and *NSt* lines retain the ability of the Tam3 TPase to recognize and bind the Tam3 sequence because EMSA showed the ability of nuclear Tam3 TPase to bind to the 26 bp DNA sequence containing three GCHCG motifs (Fig. 3C-D). Thus, it is most likely that both *OSt* and *NSt* alter the recognition by the Tam3 TPase of its binding motifs. In general, DNA methylation and histone modification could create barriers on the DNA sequence to prevent recognition and binding of DNA-binding proteins, resulting in transcriptional inhibition (Slotkin and Martienssen, 2007; Miller and Grant, 2013; Liyanage et al., 2014). However, our results do not support the involvement of DNA methylation or histone modification, as the transcription of *Tam3 TPase* occurs normally (Supplementary Fig. 8). Consequently, there are two possible ways that entities derived from the *OSt* and *NSt* loci restrict the binding of Tam3 TPase to Tam3 sequences (Fig. 6). One is that the recognition sites of Tam3 TPase are covered by such entities (Fig. 6A), and the other is that such entities compete for Tam3 TPase preventing it from binding (Fig. 6B). In this study, we found that sRNAs from the x and y regions of the Tam3 element exist in *OSt* and *NSt* lines (Fig. 4). These sRNAs might compete with the Tam3 TPase for the GCHCG motifs or interact with Tam3 TPase itself, and thus the prevent the TPase from binding, resulting in the suppression of Tam3 transposition (Fig. 6). For the above reasons, the function of these sRNAs is likely different from known TEs-derived sRNAs (Okamoto and Hirochika, 2001).

**Fig. 6.**
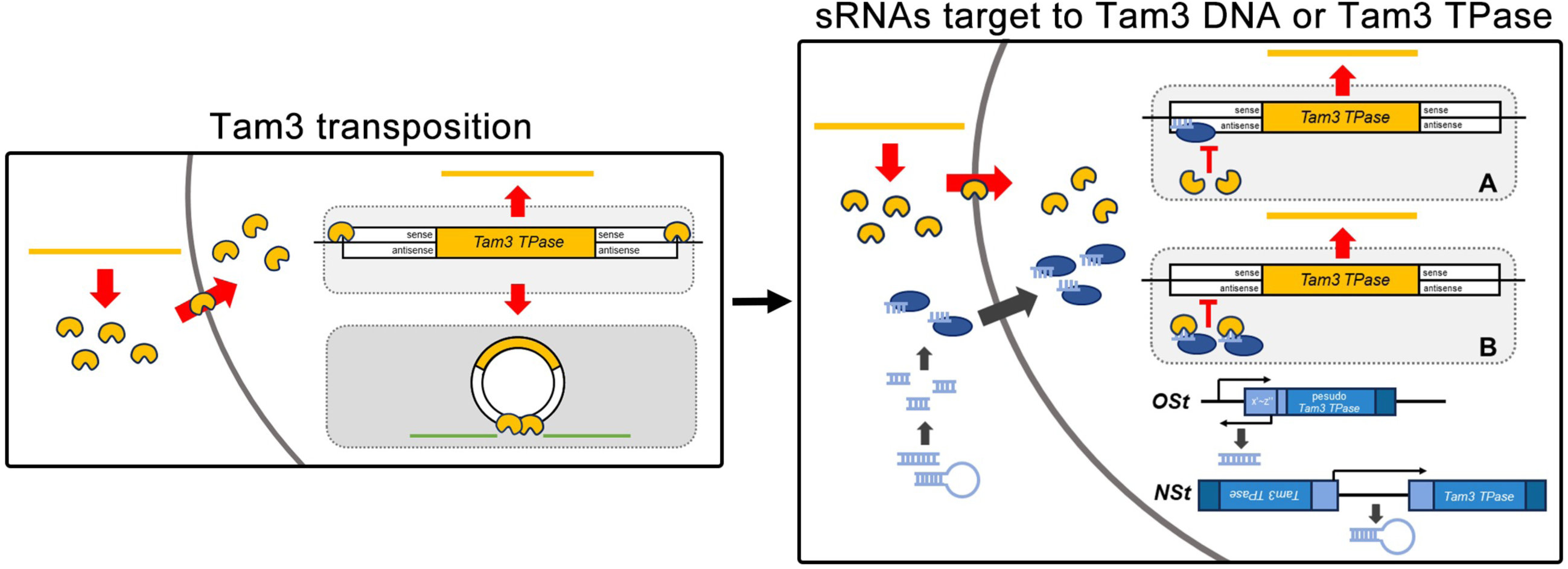
*OSt* and *NSt* suppress Tam3 transposition by two possible mechanisms. Transcription of *OSt* and *NSt* DNA fragments can result in the production of double-stranded RNAs which can be processed into sRNAs. Certain proteins carry these sRNAs and transport them into nucleus. In the nucleus, the sRNAs may bind to the antisense strand of 5’ end of Tam3 DNA (A) or interact with Tam3 TPase itself (B) to inhibit the binding between Tam3 TPase and Tam3 DNA, which results in the suppression of Tam3 transposition.

*OSt* and *NSt* could generate sRNAs that cover the target sites of Tam3 TPase (Fig. 6A) or that interact with Tam3 TPase (Fig. 6B) so that Tam3 transposition is suppressed. In the 5’ end region of the Tam3 sequence, we observed that the sRNA accumulations in the *OSt* and *NSt* lines are clearly different from those in the *ost*/*nst* line (Fig. 4). In the sense strand, there are six GCHCG motifs in the region where abundant sRNAs were observed in *OSt* and *NSt* relative to *ost*/*nst* (Fig. 4). The sRNAs accumulate in the common 5’ end region of Tam3 in the two lines suggesting that *OSt* and *NSt* suppress Tam3 transposition by similar mechanisms. However, *OSt* and *NSt* differ in the localizations of their sRNAs and in the sRNA types from the sense strand of the 5’ end of Tam3 where the Tam3 TPase binds (Fig. 4 and Supplementary Figs. 15-16). In plants, sRNAs are commonly generated from dsRNAs derived from hairpin RNAs or pairing of sense and antisense RNAs by cleavage by dicer proteins (Mathieu and Bender, 2004; Ciechanowska et al., 2021; Kryovrysanaki et al., 2022). The sRNA types preferentially detected in the *OSt* and *NSt* lines were 24 nt and 21-22 nt sRNAs, respectively. These differences could indicate that different dicer proteins function predominantly in the two lines (Supplementary Figs. 15-16). The dicer proteins are known as DICER-LIKE (DCL), and include at least DCL1, 2, 3 and 4 (Kryovrysanaki et al., 2022). Four DCL proteins are responsible for the generation of different sized sRNAs: DCL1 and 4 for 21 nt sRNAs, DCL2 for 22 nt sRNAs and DCL3 for 24 nt sRNAs (Erdmann and Picard, 2020; Kryovrysanaki et al., 2022). Normally, 24 nt sRNAs are associated with transcriptional gene silencing, whereas 21 and 22 nt sRNAs are involved in post-transcriptional gene silencing (Erdmann and Picard, 2020; Kryovrysanaki et al., 2022). It may be that different DCL proteins led to distinct sRNAs in the *NSt* and *OSt* lines, associated with their different respective precursors: double-stranded duplex in *OSt* and single RNA molecule with hairpin shape in *NSt* (Fig. 5). These results are clearly consistent with most 24 nt sRNAs in *OSt* being generated from double-stranded duplexes by DCL3, and the 21-22 nt sRNAs in *NSt* being derived from the double-stranded stem region of a hairpin-shaped RNA by DCL1 and 2 (Liu et al., 2014). These different mechanisms forming different sRNAs in *OSt* and *NSt* could be associated with semi-dominant and dominant suppressive effects of *OSt* and *NSt* respectively on Tam3 transposition, implying that the suppression by *NSt* involving the hairpin and DCL1 and 2 activity is stronger than that invoked by *OSt*.

Most TEs are inactive and are tightly regulated by hosts via master regulators, which maintain the integrity and stability of host genomes (Levin and Moran, 2011; Bhat et al., 2022). In particular, master regulators prevent large-scale TE transpositions. However, a very small proportion of TEs are active and continue to transpose under repressive conditions (Slotkin and Martienssen, 2007). The transposition of these active TEs may be regulated by more specialized mechanisms. Our work shows that TE-specific repressor genes may involve spontaneous changes that arise as a result of transposition of active TEs, resulting in self-regulation. TE-specific repressor genes can be found and detected for DNA TEs whose transposition activity can be monitored easily. Although six TE-specific repressor loci, including *OSt* and *NSt*, have now been isolated and characterized in different organisms, these loci may arise only by chance. Such self-regulatory systems could provide a means for active TEs to maintain a balanced relationship with their hosts when they escape from master regulation. Within TE self-regulatory systems, TE-specific repressor loci could be a characteristic of active TEs, at least class II, DNA TEs.

### Materials and Methods Plant Materials

Four lines of *Antirrhinum majus* were used in this study, two Tam3-active and two Tam3-repressed lines (Supplementary Fig. 2). One of the Tam3-active lines derived from the John Innes Center (JIC) stock JI: 98, here named HAM2, has the genotype for *ost*/*nst*/*niv*^rec^/*Pal* (Supplementary Fig. 2A). The other Tam3-active line called HAM5 contains the genotype for *ost*/*nst*/*niv*^rec: beni^/*Pal* (Supplementary Fig. 2B). This line was isolated from the progeny of JI: 98 (H101) carrying the *niv*^rec^ allele crossed with the two different lines: Stock 44 and 45 in the JIC collections (Uchiyama et al., 2013). In the two Tam3-repressed lines: the JIC stock JI: 558 carrying *OSt* had been produced by crossing *niv^rec^*:98 to the original *Stabiliser* line, stock JI: 3 (*OSt*/*nst*/*Niv*/*pal*^rec^) (Harrison and Carpenter, 1973), and the *OSt*/*nst*/*niv*^rec^/*Pal* genotype had been selected, here named HAM21. The other *St* line, called HAM3, carrying *ost*/*NSt*/*niv*^rec: beni^/*Pal*, which is an isogenic line of HAM5 (o*st*/*nst*/*niv*^rec: beni^/*Pal*) (Uchiyama et al., 2008). To supplement the *OSt*/*niv*^rec^ (HAM21) and *ost*/*niv*^rec^ (HAM2) test cross, we also crossed JIC Stock2 (*ost*/*nst*/*Niv*/*pal*^rec^) named HAM22, to HAM21 and examined the segregation of the petal variegation trait in the F_2_ population. To obtain F_2_ seeds, we made three combinations of crosses [*ost*/*niv*^rec^ (HAM2) × *OSt*/*niv*^rec^ (HAM21)], [*ost*/*Niv*/*pal*^rec^ (HAM22) × *OSt*/*niv*^rec^/*Pal* (HAM21)], and [*nst*/*niv*^rec: beni^ (HAM5) × *NSt*/*niv*^rec: beni^ (HAM3)]. Plants were initially grown at 25℃ for 1.5 months and transferred to 15℃ or 25℃ growth chambers and grown for at least 2 weeks to score their petal variegation phenotypes.

### DNA extraction and PCR analysis

DNA was extracted from young leaves (3-4 cm in length) of *Antirrhinum* plants using extraction buffer (100 mM Tris-HCl [pH 8.0], 10 mM EDTA [pH 8.0], 1 M KCl) combined with phenol/chloroform extraction method. PCR analysis was performed using genomic DNA. The PCR solution containing 200 ng of DNA and components of Ex Taq DNA polymerase (Takara Bio, Shiga Japan) was denatured at 94 ℃ for 30 s, annealed at appropriate temperatures based on the melting temperature for each primer for 30 s, and extended at 72 ℃ for a given time based on the length of the amplified fragments for a total of 30 cycles. Each primer set used for PCR was listed in Supplementary Table 4.

### Phenotypic segregations in the F_2_ lines and transposon tagging

Based on flower variegation phenotypes at the *Niv* locus, phenotypic segregations were investigated using F_2_ plants obtained by selfing F_1_ plants from crosses between the highly variegated lines [*ost*/*niv*^rec^ (HAM2) or *nst*/*niv*^rec: beni^ (HAM5)] and the *Stabiliser* lines *NSt/niv*^rec: beni^ (HAM3) or *OSt/niv*^rec^ (HAM21). The F_2_ plants between *ost*/*niv*^rec^ (HAM2) and *OSt*/*niv*^rec^ (HAM21) were sorted into high- (>2500 spots per flower likely *ost*/*ost; niv*^rec^/*niv*^rec^), middle- (100-500 spots per flower likely *OSt*/*ost; niv*^rec^/*niv*^rec^) and low- (<20 spots per flower likely *OSt*/*OSt; niv*^rec^/*niv*^rec^) spotted plants and were used for transposon tagging (Supplementary Fig. 2A). For *NSt*, F_2_ plants between *nst*/*niv*^rec: beni^ (HAM5) and *NSt*/*niv*^rec: beni^ (HAM3) were segregated into high spot (> 1000 spots per flower likely *nst*/*nst; niv*^rec: beni^/*niv*^rec: beni^) and low spot (<10 spots per flower, likely *NSt*/*NSt; niv*^rec: beni^/*niv*^rec: beni^ and *NSt*/*nst; niv*^rec: beni^/*niv*^rec: beni^) (Supplementary Fig. 2B).

A possible reason for the *Stabiliser* phenotypes has long been proposed to be related to alterations in Tam3 itself (Fincham and Harrison, 1967). To investigate the relationship between Tam3 and petal variegation phenotypes, Southern hybridizations were performed for both F_2_ segregations in the *OSt* and *NSt* populations using the Tam3 sequence as a probe. For Southern hybridizations, *Eco*RI- and *Eco*RV-digested genomic DNAs for the *OSt* and *NSt* populations, respectively, were transferred to Hybond-N+ membranes (Pall, http://www.pall.com); all procedures were performed using an ECL Direct Nucleic Acid Labeling and Detection System in accordance with the manufacturer’s instructions (GE Healthcare, http://www.gelifesciences.co). The procedures of Tam3 transposon tagging for *OSt* and *NSt* are detailed in Supplementary Description 1 and 2.

In the Southern hybridization pattern of the *OSt* F_2_ population, we found the 6.6 kb-Tam3 *Eco*RI fragment linked to the *St* phenotypes (Supplementary Fig. 4A), while the hybridization pattern in the *NSt* F_2_ population showed 3.5 kb-Tam3 *EcoR*V fragment linked to the *St* phenotypes (Supplementary Fig. 6A). To identify the *St*-linked DNA fragment from each of the *OSt* and *NSt* populations, we followed the Southern hybridization methods described by Kishima et al. (1999). The bands recovered from the gels after the electrophoresis at the appropriate sizes were cloned into a lambda phage vector (Uni-ZAP XR vector: Agilent Technologies) or plasmid vector (pBluescriptSK+: Agilent Technologies) and screened by plaque or colony hybridization using the entire Tam3 sequence as a probe. Selected clones were sequenced by the Sanger method. The association with the *OSt*/*ost* phenotypes was validated for 64 individuals by two PCR amplifications: one primer combination of R [ATA AAC CCT AAC GAA CGA GGT GGA AAT G] and F1 [GAA CCC CAC TCG AAG ACG GAT TG] amplified the sequence without the *OSt*-Tam3 insertion for the *ost* allele (3.0 kb), and the other primer combination of R and F2 [CCA ACA ATC CAA CCA ACC AAG] resulted in amplification of the sequence with the *OSt*-Tam3 insertion for the *OSt* allele (1.5 kb) (Supplementary Fig. 4C; Supplementary Table 1). The linkage of the *NSt* locus with the tagged sequence was examined and confirmed for more than 200 individuals by PCR with two primer combinations: one primer combination of F1 [AAG GCT TTC ATT GCA TAG GGA TCT GG] and R1 [CCA AGT ACT GAA TAA GCA GGA GGT GAA] amplified the sequence without the Tam3-R insertion for *nst* allele (0.7 kb), and the other combination of F2 [GAA CCG GCT GGA GAA ATG] and R2 [AGC TGC AGC ACA AAA AAC AGG CCT ATA C] gave rise to the amplification of the sequence with Tam3-R insertion for *NSt* allele (0.33 kb), and Southern hybridization with the probe of the flanking sequence to Tam3 in the selected clone (Supplementary Fig. 6C and D; Supplementary Table 1). The chromosomal positions for the *OSt* and *NSt* loci were determined with Snapdragon Genome Database at http://bioinfo.sibs.ac.cn/Am/index.php.

### RNA extraction and qRT-PCR

Total RNA was extracted from young leaves of *Antirrhinum* plants using PureLink^®^ RNA Mini Kit (Thermo Fisher Scientific). Approximately 800 ng of total RNA was reverse transcribed into cDNA using QuantiTect Reverse Transcription Kit (QIAGEN) according to the manufacturer’s instructions. qRT-PCR was performed to detect the expression level of *Tam3 TPase* gene on CFX Connect^TM^ Real-Time System (BIO-RAD) using SYBR^®^ GreenER^TM^ qPCR SuperMix (Thermo Fisher Scientific). The *Antirrhinum UBIQ* gene was used as internal control. Primers used for qRT-PCR are listed in Supplementary Table 4.

### Total protein extraction

The leaf powder of *Antirrhinum* plant material was mixed with the protein extraction buffer (50 mM Tris-HCl [pH 7.5], 150 mM NaCl, 1 mM EDTA, 10% glycerol, 5 mM DTT, 0.2% TritonX-100, and protease inhibitor cocktail [Sigma-Aldrich]), and the homogenate was incubated on ice for 30 min with shaking. After incubation, the homogenate was centrifuged at 13,000 rpm for 15 min at 4 ℃. The supernatant was collected and centrifuged at 13,000 rpm for 10 min at 4 ℃, and the resulting supernatant was collected as total protein. The protein concentration was determined using a protein assay kit (Bio-Rad) before use.

### Nuclear protein extraction

At least 1.0 g of *Antirrhinum* plant leaf powder was thoroughly mixed with Honda buffer (0.44 M sucrose, 20 mM HEPES-KOH [pH 7.4], 10 mM MgCl_2_, 1 mM DTT, 1.25% Ficoll, 2.5% dextran T40, 0.5% TritonX-100, and protease inhibitor cocktail [Sigma-Aldrich]), and the homogenate was filtered through two layers of Miracloth (Sigma-Aldrich). The filtered homogenate was centrifuged at 3,500 rpm for 5 min at 4 ℃, and the supernatant was removed. The nuclear pellet was resuspended in Honda buffer and centrifuged at 3,500 rpm for 5 min at 4℃. The washing step was repeated for three times, and the final white nuclear pellet was resuspended in protein extraction buffer (50 mM Tris-HCl [pH 7.5], 150 mM NaCl, 1 mM EDTA, 10% glycerol, 5 mM DTT, 0.2% TritonX-100, and protease inhibitor cocktail [Sigma-Aldrich]). The nuclear mixture was incubated on ice for 30 min with shaking, then centrifuged at 15, 000 × g for 20 min at 4℃. After centrifugation, the supernatant was collected as nuclear protein. The protein concentration was determined using a protein assay kit (Bio-Rad) before use.

### Western blot analysis

Protein extracts were mixed with 5˟ protein loading buffer and denatured at 100 ℃ for 10 min. Denatured protein was separated on 10% SDS-PAGE gel and transferred to a polyvinylidene fluoride (PVDF) membrane. After blocking with 5% (w/v) skim milk at 4 ℃ overnight, the membrane was incubated with primary antibody for 1 hour at room temperature. The membrane was incubated with 5000-fold diluted secondary antibody (goat anti-rabbit IgG [H+L] horseradish peroxidase conjugate, BIO-RAD) for 1 hour at room temperature and stained with ECL detection reagents (GE Healthcare) according to the manufacturer’s instructions. Signals were detected using a Fujifilm LAS4000mini Luminescence Imaging Analyzer.

### Transient transformation of protoplast and microscopic observation

Protoplasts were isolated from young *Antirrhinum* leaves and transiently transformed with approximately 15 ug of pA7-Tam3 TPase-GFP plasmid DNA, used in Zhou *et al*. (2017), using PEG-calcium transfection method according to Yoo *et al*. (2007). After transient transformation, all samples were cultured at 25 ℃ for 20 h. Upon completion of culture, fluorescence signals were observed using an all-in-one fluorescence microscope (BZ-X810, Keyence).

### EMSA

Nuclear protein was extracted as described above. The DNA probe used in Hashida *et al*. (2006) contained three Tam3 TPase binding motifs (GCHCG), and its nucleotide position is from 79 to 104 in the Tam3 sequence (see in Fig. 3D). To synthesize the DNA probe, the biotin-labelled positive-strand oligonucleotide was equally mixed with the negative-strand oligonucleotide, and the oligonucleotide mixture was incubated at 95℃ for 10 min and cooled down to produce the DNA probe. EMSA was performed using the LightShift ^®^ chemiluminescent EMSA kit according to the manufacturer’s instructions.

### RNA-seq analyses

Total RNA was extracted as described above. gDNA was removed from RNA using Dnase I (Nippon Gene). RNA library preparation and sequencing were performed by Rhelixa (Japan), and 150 bp paired-end (PE150) reads were generated.

For data analysis, raw sequences were processed to remove adapters and low quality sequences using Trimmomatic (Bolger et al., 2014). Quality control of trimmed reads was performed using FASTQC (https://www.bioinformatics.babraham.ac.uk/projects/fastqc/). The trimmed reads were then aligned to reference sequences using hisat2 (http://daehwankimlab.github.io/hisat2/). Aligned SAM files were converted to BAM files using Samtools (https://www.htslib.org/), then read counts were performed on BAM files using Samtools as well.

### sRNA-seq analyses

Total RNA was extracted as described above. gDNA was removed from RNA using Dnase I (Nippon Gene). sRNA library and sequencing were performed by Rhelixa (Japan).

For the data analysis, raw reads were processed to remove adapters and low-quality reads using Trimmomatic (Bolger et al., 2014). In trimmed reads, reads that contain ≥ 20 A or whose length is < 15 nt or > 35 nt were removed using cutadapt (https://cutadapt.readthedocs.io/en/stable/). After read cutting, quality control of the reads was carried out using FASTAQC (https://www.bioinformatics.babraham.ac.uk/projects/fastqc/). Then, rRNAs, tRNAs, snRNAs, snoRNAs and known miRNAs were removed from qualified reads using Rfam (https://rfam.org/) and miRbase (https://mirbase.org/) databases. The processed reads were aligned to reference sequences using bowtie2 (https://bowtie-bio.sourceforge.net/bowtie2/index.shtml). Aligned SAM files were converted to BAM files using Samtools (https://www.htslib.org/). The screening of perfectly aligned reads and three sized sRNAs (21, 22 and 24 nt), and read counting were carried out with BAM files using Samtools.

### Data depositions

The genomic sequences of the regions including *OSt* and *NSt* were deposited in NCBI GenBank PP898194 and PP898195, respectively. The RNA sequencing data for mRNA, long non-coding RNA (lncRNA), and small RNA (sRNA) were registered in NCBI Sequence Read Archive SAMN41273101 (HAM3 mRNA), SAMN41273102 (HAM21 mRNA), SAMN41273103 (HAM5 mRNA), SAMN41273104 (HAM3 lncRNA), SAMN41273105 (HAM21 lncRNA), SAMN41273106 (HAM5 lncRNA), SAMN41273107 (HAM3 sRNA), SAMN41273108 (HAM21 sRNA), and SAMN41273109 (HAM5 sRNA).

## Acknowledgements

We gratefully acknowledge Dr. Y. Koide (Hokkaido University) for his valuable suggestions and comments. We also thank H. Zhou, S. Mitsui, H. Asano, and R. Tanabe (Hokkaido University) for technical assistances. Seeds were kindly supplied by R. M. Carpenter and L. Copsey (John Innes Centre). This work was supported by JSPS KAKENHI grant #08760002.

## Author contributions

A. C. M. and Y.K. made the initial concept of the study. W.S., T.U., and Y.K. designed the experimental plan. W.S., C.M., and Y.K. drafted the manuscript with input from all other authors. Y.K., T.U. and H.K. performed the transposon tagging. S.W., M.H., and I.Y. analyzed and evaluated genetic traits and phenotypes. S.W. and K.N. performed immunoblotting analyses and EMSA. S.W. performed most of the other molecular analyses.

## Abbreviations

*Stabiliser* (*St*), *Old Stabiliser* (*OSt*), *New Stabiliser* (*NSt*), Tam3 transposase (Tam3 TPase), small RNAs (sRNAs), transposable element (TE), master regulators (MRs), *Palllida* (*pal*), *Nivea* (*niv*), BED zinc finger motif (Znf-BED), open reading frame (ORF), long non-coding RNA (lncRNA), DICER-LIKE (DCL).

---

**Supplementary Fig. 1**

Profile of wild type (WT) *A. majus* line.

WT contains the *Niv* allele at *Niv* locus, where no Tam3 is inserted, and thus the flower petals exhibit red color at 25℃ and 15℃. Additionally, the observation of 2.0 kb bands at 25℃ and 15℃ indicates no Tam3 insertion in the *Niv* allele.

**Supplementary Fig. 2**

F2 segregation profiles in the *OSt*/*niv*^rec^ × *ost*/*niv*^rec^ (HAM21×HAM2) and *NSt*/*niv*^rec: beni^ × *nst*/*niv*^rec: beni^ (HAM3×HAM5) *Antirrhinum* crossed lines.

A. F2 segregation in the *OSt*/*niv*^rec^ × *ost*/*niv*^rec^ (HAM21×HAM2) cross. C. F2 segregation in the *NSt*/*niv*^rec: beni^ × *nst*/*niv*^rec: beni^ (HAM3×HAM5) cross. The number of plants showing different spotted phenotypes in F2 is shown in Supplementary Table 1.

**Supplementary Fig. 3**

Chromosomal positions of *OSt*, *NSt*, *Niv* and *Pal* within the *Antirrhinum majus* genome.

**Supplementary Fig. 4**

Identification of *OSt*.

A.Transposon tagging analysis in *ost*/*niv*^rec^ (HAM2), *OSt*/*niv*^rec^ (HAM21) and their progeny plants. Each DNA was extracted and digested with *Eco*RI. The ORF region of Tam3 was used as a probe, and black arrow indicates the 6.6 kb “tag” bands in *OSt*/*niv*^rec^ (HAM21), F1 and middle-spotted F2 plants. B. Different flower variegation phenotypes of plants used for the identification of *OSt*. C. Structures of *OSt* and *ost* alleles. The two thin vertical bars in *OSt* represent *Eco*RI sites. Arrows represent the positions of primers present in *OSt* and *ost* alleles. D. PCR analysis of the allelic constitution at *OSt* locus in plants with different flower petal variegation phenotypes. The A (1.5 kb) fragment amplified using the F2 and R primers indicates the *OSt* allele, while the B (3.0 kb) fragment amplified using the F1 and R primers indicates the *ost* allele.

**Supplementary Fig. 5**

Dot plot analysis and sequence alignment of the *OSt* sequence.

A. *OSt* sequence was aligned to the intact Tam3 sequence from *niv*^rec^. In the dot plot, the upper horizontal axis represents *OSt* sequence, and left vertical axis represents the Tam3 sequence. Red and green dots represent the alignment of the sequence of positive and negative strands of *OSt* to the sequence of the positive strand of Tam3, respectively. The window size and mismatch limit settings in dot plot analysis are 10 and 0, respectively. B. Sequence alignment between x-y-z region of intact Tam3 and x’-y’-z’-x’’-z’’ region of *OSt*-Tam3. Black and blue squares indicate SNP and deletion positions, respectively. Red and black underlines indicate Terminal Inverted Repeat (TIR) and GCHCG motifs, respectively.

**Supplementary Fig. 6**

Identification of *NSt*.

A. Transposon tagging analysis in low- and high-spotted plants. Each DNA was extracted and digested with *Eco*RV. Full-length Tam3 sequence was used as a probe, and black arrow indicated 3.5 kb tagging bands in low-spotted plants. B. Different flower variegation phenotypes of plants used for the identification of *NSt*. C. Structures of *NSt* and *nst* alleles. Vertical bars indicate the nucleotide positions of restriction enzymes including *Eco*RV, *Bgl*I and *Xba*I in *NSt* and *nst* alleles. The bold black line below *nst* in the diagram represents the probe used for Southern blotting analysis. D. Southern blotting analysis of allelic constitution at *NSt* locus in plants with different flower petal variegation phenotypes. Each DNA was extracted and digested with *Eco*RV. The probe used is shown in the C diagram. The 3.5 kb fragment was detected in low-spotted plants and indicates the *NSt* allele, while the 2.7 kb fragment was detected in high-spotted plants and indicates the *nst* allele.

**Supplementary Fig. 7**

Dot plot analysis of the *NSt* sequence and the process of *NSt* formation.

A. *NSt* sequence was aligned to the intact Tam3 sequence from *niv*^rec^. In the dot plot, the upper horizontal axis represents *NSt* sequence, and the left vertical axis represents the Tam3 sequence. Red and green dots represent the alignment of the sequence of positive and negative strands of *OSt* to the sequence of positive strand of Tam3, respectively. The window size and mismatch limit settings for the dot plot analysis were 10 and 0, respectively. B. Proposed process of *NSt* formation. One Tam3 element was inserted into the third intron of *Am02g32060* gene, which results in *nst* formation. *NSt* formation was due to the insertion of second Tam3 element on the right side of first Tam3 element in the same intron.

**Supplementary Fig. 8**

The transcription and translation of *Tam3 TPase*.

A. qRT-PCR analysis of *Tam3 TPase* expression. Expression levels were normalized to that of *UBIQ* (X67957). Data represent the means ± SD (n = 3). Significant differences were determined by Student’s t-test. The experiments were performed three times. B. Immunoblot analysis of Tam3 TPase in the same plants employed with the qRT-PCR analysis. Total protein was extracted from young leaves of *Antirrhinum* plants. The fragments indicated by arrows correspond to 100 kDa of Tam3 TPase.

**Supplementary Fig. 9**

Distributions and amounts of mRNAs in Tam3 region.

The mRNAs that were perfectly matched to the Tam3 region (1-3633 bp) were analyzed. Red and blue colors were used to distinguish sense and antisense mRNAs, respectively. The vertical fold arrow in the Tam3 structure diagram represents the transcriptional start site of *Tam3 TPase* gene. ATG and TAA are start and stop codons of Tam3 TPase, respectively. FPH, fragments per hundred mapped fragments.

**Supplementary Fig. 10**

Distributions and amounts of mRNAs in Tam3 and *OSt*-Tam3 regions in *OSt*/*niv*^rec^ (HAM21).

The mRNAs that perfectly matched were aligned to the Tam3 region (1-3633 bp) and the *OSt-*Tam3 region (1-3919 bp) in *OSt*/*niv*^rec^ (HAM21). Red and blue colors were used to distinguish sense and antisense mRNAs, respectively. The vertical fold arrow in the Tam3 structure diagram represents the transcriptional start site of the *Tam3 TPase* gene. ATG and TAA are start and stop codons of Tam3 TPase, respectively. FPH, fragments per hundred mapped fragments.

**Supplementary Fig. 11**

Distributions and amounts of mRNAs in *NSt* and *nst* alleles.

The mRNAs that perfectly matched were aligned to the 5’ region of Tam3-L of *nst* (1-689 bp) and the flanking region of Tam3-L of *nst* (1867 bp) in *nst*/*niv*^rec: beni^ (HAM5), and aligned to 5’ regions of Tam3-L (1-689 bp) and Tam3-R (1-689 bp) of *NSt* and internal region (1867 bp) between Tam3-L and Tam3-R of *NSt* in *NSt*/*niv*^rec: beni^ (HAM3). Red and blue colors were used to distinguish sense and antisense mRNAs, respectively. FPH, fragments per hundred mapped fragments.

**Supplementary Fig. 12**

Distributions and amounts of RNAs (lncRNAs + mRNAs) in the Tam3 region.

The RNAs including lncRNAs and mRNAs that were perfectly matched to the Tam3 region (1-3633 bp) were analyzed. Red and blue colors were used to distinguish sense and antisense RNAs, respectively. The vertical fold arrow in the Tam3 structure diagram represents the transcriptional start site of the *Tam3 TPase* gene. ATG and TAA are start and stop codons of the Tam3 TPase, respectively. FPH, fragments per hundred mapped fragments.

**Supplementary Fig. 13**

Distributions and amounts of RNAs (lncRNAs + mRNAs) in Tam3 and *OSt*-Tam3 regions in *OSt*/*niv*^rec^ (HAM21).

The RNAs including lncRNAs and mRNAs that perfectly matched were aligned to the Tam3 region (1-3633 bp) and the *OSt*-Tam3 region (1-3919 bp) in *OSt*/*niv*^rec^ (HAM21). Red and blue colors were used to distinguish sense and antisense RNAs, respectively. The vertical fold arrow in the Tam3 structure diagram represents the transcriptional start site of the *Tam3 TPase* gene. ATG and TAA are start and stop codons of the Tam3 TPase, respectively. FPH, fragments per hundred mapped fragments.

**Supplementary Fig. 14**

Distributions and amounts of RNAs (lncRNAs + mRNAs) in *NSt* and *nst* alleles.

The RNAs including lncRNAs and mRNAs that perfectly matched were aligned to the 5’ region of Tam3-L of *nst* (1-689 bp) and the flanking region of Tam3-L of *nst* (1867 bp) in *nst*/*niv*^rec: beni^ (HAM5), and aligned to the 5’ regions of Tam3-L (1-689 bp) and Tam3-R (1-689 bp) of *NSt* and the internal region (1867 bp) between Tam3-L and Tam3-R of *NSt* in *NSt*/*niv*^rec: beni^ (HAM3). Red and blue colors were used to distinguish sense and antisense RNAs, respectively. FPH, fragments per hundred mapped fragments.

**Supplementary Fig. 15**

Distributions and amounts of sense sRNAs in the Tam3 region.

Except for *OSt*, sRNAs that were perfectly matched to the sense strand of Tam3 (1-3633 bp) were analyzed in *NSt* and *ost*/*nst*. In *OSt*, sRNAs that contained zero and one SNP were aligned to the sense strand in the 5’ 1-201 bp region of Tam3, and sRNAs were perfectly aligned to the sense strand in 202-3633 bp region of Tam3. The vertical fold arrow and red vertical line in the Tam3 structure diagram represent the transcriptional start site of the *Tam3 TPase* gene and the position of the GCHCG motifs, respectively. ATG and TAA are start and stop codons of the Tam3 TPase, respectively. RPH, reads per hundred mapped reads.

**Supplementary Fig. 16**

Distributions and amounts of antisense sRNAs in the Tam3 region.

Except for *OSt*, sRNAs that were perfectly matched to the antisense strand of Tam3 (1-3633 bp) were analyzed in *NSt* and *ost*/*nst*. In *OSt*, sRNAs that contained zero and one SNP were aligned to the antisense strand in the 5’ 1-201 bp region of Tam3, and sRNAs were perfectly aligned to the antisense strand in the 202-3633 bp region of Tam3. The vertical fold arrow and red vertical line in the Tam3 structure diagram represent the transcriptional start site of the *Tam3 TPase* gene and the positions of GCHCG motifs, respectively. ATG and TAA are start and stop codons of the Tam3 TPase, respectively. RPH, reads per hundred mapped reads.

**Supplementary Fig. 17**

Distributions and relative amounts of total sRNAs in 3’ region of intact Tam3.

sRNAs that were perfectly matched to the 3’ region of Tam3 (3433-3633 bp) were analyzed. Red and blue squares on nucleotide sequences indicate Terminal Inverted Repeat (TIR) and GCHCG motifs, respectively. Lines indicate as follows; blue: *OSt* (HAM21), red: *NSt* (HAM3), black: *ost*/*nst* (HAM5). Relative ratios were calculated as follows; the RPH (reads per hundred mapped reads) of each line divided by the average RPH of *ost*/*nst* line within 3’ 3433-3633 bp region of Tam3.

**Supplementary Fig. 18**

Distributions and amounts of sRNAs in *OSt-*Tam3 region.

sRNAs that were perfectly matched to the *OSt-*Tam3 region (1-3919 bp) were analyzed. Red and blue colors were used to distinguish sense and antisense sRNAs, respectively. RPH, reads per hundred mapped reads.

**Supplementary Fig. 19**

Distributions and amounts of sRNAs in 5’ regions of Tam3, *OS*t-Tam3 and Tam3 of *nst* and *NSt*.

A. Distributions and amounts of sRNAs in 5’ x-y-z region of Tam3 in *nst*/*niv*^rec: beni^ (HAM5). B. Distributions and amounts of sRNAs in 5’ x’-y’-z’-x’’-z’’ region of *OSt*-Tam3 in *OSt*/*niv*^rec^ (HAM21). C. Distributions and amounts of sRNAs in the 5’ region of Tam3-L of *nst* (660-689 bp) and flanking region of Tam3-L of *nst* (690-740 bp) in *nst*/*niv*^rec: beni^ (HAM5). D. Distributions and amounts of sRNAs in the 5’ regions of Tam3-L (660-689 bp) and Tam3-R (2557-2585 bp) of *NSt* and flanking regions of Tam3-L (690-740 bp) and Tam3-R (2503-2556 bp) in *NSt*/*niv*^rec: beni^ (HAM3). sRNAs that perfectly matched were aligned to the regions. Gray, red squares on nucleotide sequences indicate Target Site Duplication (TSD), Terminal Inverted Repeat (TIR), respectively. Lines indicate as follows; blue: total sRNAs, green: 24nt sRNAs, black: 22nt sRNAs and red: 21nt sRNAs. RPH, reads per hundred mapped reads.

**Supplementary Fig. 20**

Detection of a possible hairpin transcript in the *NSt* and *nst* alleles.

In panel A, the F1 and R1 primers amplified a 0.8 kb PCR fragment in the DNA and cDNA samples with the *NSt* allele and DNA sample with *nst* allele, but not in the RNA direct sample with the *NSt* allele and RNA samples with the *nst* allele. Panel B shows a 1.5 kb PCR fragment amplified with the DNA and cDNA of *NSt* and *nst* alleles using F2 and R2 primers. F1 and R1 are located at the 5’ end of both Tam3-L and Tam3-R. F2 and R2 primers are presented in the internal region between two Tam3 elements of *NSt* (1-1867 bp). Supplementary Table 4 shows the primer sequences.

